# *Bacillus subtilis* with a 1% population enhances the activity of *Enterococcus faecalis* with a 99% population

**DOI:** 10.1101/2021.09.30.462685

**Authors:** Tsukasa Ito, Yu Yamanashi

**Affiliations:** Department of Environmental Engineering Science, Gunma University, Kiryu, Gunma, Japan

**Keywords:** microbial interaction, dye decolorization assay, minority, redox potential (ORP), amino acids, metabolomics

## Abstract

Microbes are present as communities in the environment. However, the importance of minor populations has not been well studied experimentally. In this study, we evaluated the role of *Bacillus subtilis* with a 1% population and its effect on co-incubated *Enterococcus faecalis* with a 99% population. Here we used an azo dye-decolorizing *Enterococcus faecalis* strain T6a1 and non-dye-decolorizing *Bacillus subtilis* strain S4ga. The dye decolorization assay enabled the investigation of the effects of *B. subtilis* S4ga on the activity of *E. faecalis* T6a1, even when *B. subtilis* S4ga was present at only 1% relative abundance or lower. We found that non-decolorizing *B. subtilis* S4ga enhanced the dye decolorization activity of *E. faecalis* T6a1, shortened the lag time of *E. faecalis* T6a1 to start decreasing the dye concentration, and increased the time for *E. faecalis* T6a1 to continue dye decolorization. These effects were correlated with redox potential values. We compared the extracellular amino acids between each incubation culture of *E. faecalis* T6a1 and *B. subtilis* S4ga, which revealed their mutual relationship by cross-feeding of specific amino acids. We also compared the intracellular primary metabolites between co-incubation and sole incubation of *E. faecalis* T6a1. The arginine deiminase (ADI) pathway in the co-incubated *E. faecalis* T6a1 was activated compared to that of *E. faecalis* T6a1 incubated solely. These findings explained that co-incubation with *B. subtilis* S4ga promoted ATP production in *E. faecalis* T6a1 cells to a greater extent and enhanced dye-decolorization activity.

**IMPORTANCE:** This study highlights the importance of minor bacterial populations and their effects on major populations. We used *Enterococcus faecalis* as the major population and *Bacillus subtilis* as the minor population. Both species are becoming increasingly important. Some strains of *E. faecalis* are antibiotic-resistant pathogens, show probiotic effects, or are applicable for textile wastewater treatment. Some strains of *B. subtilis* are known to produce antimicrobial agents, reduce intestinal inflammation, or restore gut microbiota. We demonstrated that a low abundance of *B. subtilis* with 1% population increased the amount of energy produced by *E. faecalis* with 99% population, which appeared as enhanced dye decolorization activity of dye-decolorizing *E. faecalis*. Metabolomic analysis suggested that *E. faecalis* and *B. subtilis* had a mutual relationship by feeding specific amino acids to each other. These results provide new insights into co-existing minor populations in microbial communities and will improve our understanding of bacterial control.

**Graphical abstract:** 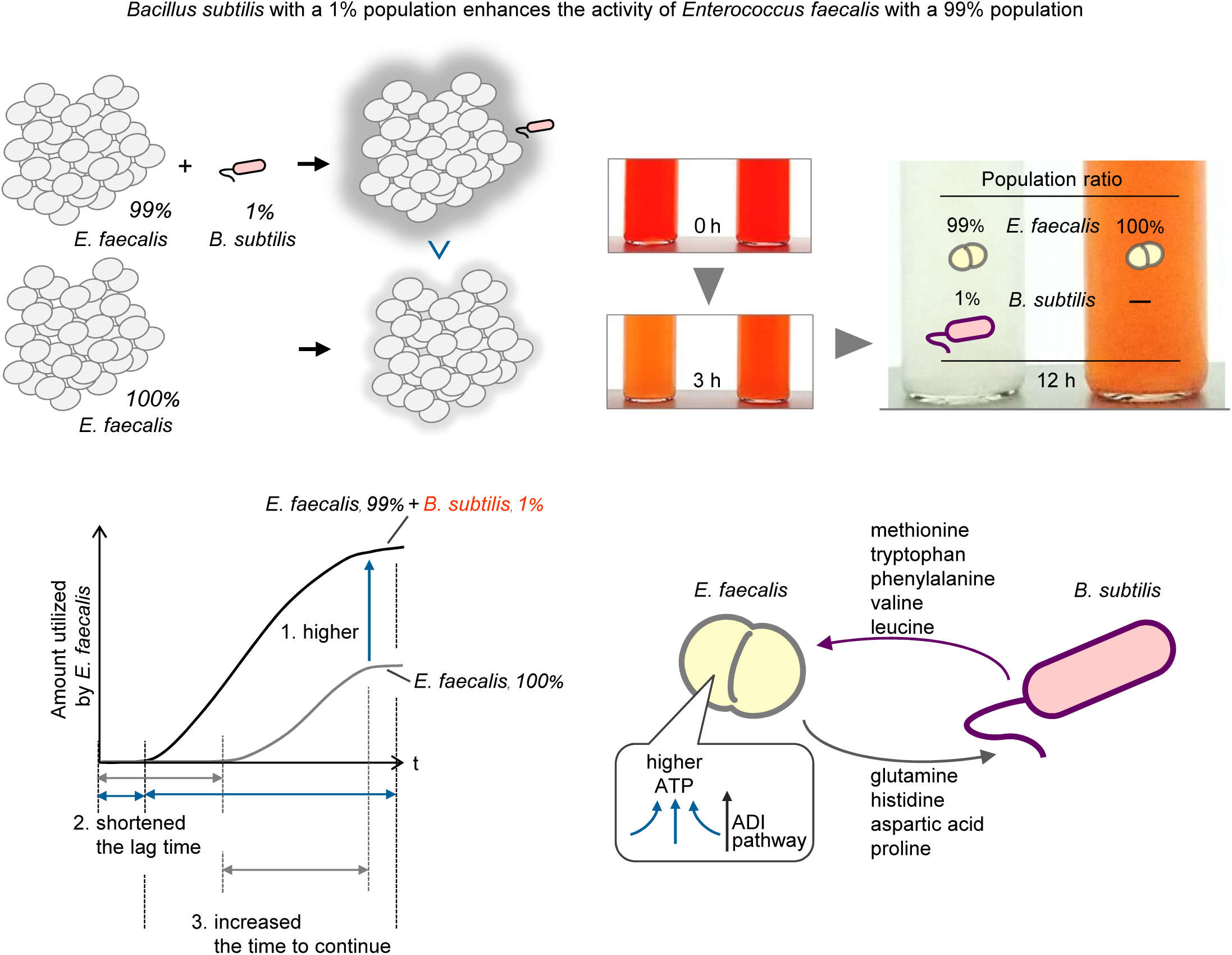

## INTRODUCTION

*Enterococcus faecalis* is becoming an enormously important bacterium in various aspects. Some strains are pathogens and are nosocomial antibiotic-resistant bacteria known as vancomycin-resistant enterococci (**1, 2**), while some strains produce an antimicrobial agent, bacteriocin, against spoilage bacteria (**3, 4**) or add flavoring (**4**) in the food industry. Probiotics for humans and animals have also been reported (**4, 5**). In the environment, *E. faecalis* is a fecal indicator and is used for microbial source tracking (**6**), and other strains are dye-decolorizing bacteria applicable for wastewater treatment discharged from the dye/textile industry (**7, 8**). Since *E. faecalis* is present in communities with other bacteria in the above situations, investigating how *E. faecalis* is associated with other bacteria is the key to establishing a way to control *E. faecalis*.

*Bacillus subtilis* is known for its beneficial effects by producing antimicrobial agents and surfactants (**9, 10**), by reducing intestinal inflammation (**9, 11, 12**), and by modulating or restoring the gut microbiota (**13, 14**), despite its lower abundance in the microbial communities. Here we focused on the combination of *B. subtilis* and *E. faecalis* and on their relationship when *B. subtilis* is a minor population, such as an additive.

Although next-generation sequencing technology currently enables the detection of microbial community members with very low abundances, their functional roles in microbial communities are poorly understood. To investigate the role of the minor population, it is necessary to establish a stable community composition during the investigation and to quantify the population composition. Furthermore, it is necessary to assay the functional activity of the major population affected by minor populations. Some assays may be required to separate or distinguish minor populations from mixed communities.

In this study, we used a dye-decolorizing *E. faecalis* strain as a reporter strain, such that when the strain is activated its decolorization is enhanced, and a non-dye-decolorizing *B. subtilis* strain as the partner strain with a minor population. We evaluated the activities of *E. faecalis* when incubated solely and when co-incubated with *B. subtilis* using the dye decolorization assay. The dye decolorization assay, though a low-tech method, visualized the activity of *E. faecalis* as the color change of test tubes within a day with high throughput. Although green fluorescent protein (GFP) has been used in many reporter assays, GFP-generating fluorophores require oxygen (**15**). The GFP-independent activity assay (dye decolorization assay) is suitable for targeting the dye-decolorizing enzyme azoreductase of *E. faecalis*, which is activated in reduced environments (**16**). Another benefit is that *E. faecalis* and *B. subtilis* are easily culturable, distinguishable, and therefore quantifiable on the same agar plate even though their population ratio is 100:1 or more. Furthermore, we were able to separate each strain from the co-incubated culture by using a 0.8-μm filter for analyzing intracellular and extracellular metabolites. In the dye decolorization assay, we prepared the initial *E. faecalis* population to be approximately 7–8 log CFU/mL and incubated in 10% Luria-Bertani (LB) medium, which controlled the growth of *E. faecalis* and *B. subtilis* during co-incubation. For example, although *B. subtilis* can grow from 5 log CFU/mL to 8 log CFU/mL within a day, it was restrained to 6 CFU/mL when co-incubated with 8 log CFU/mL of *E. faecalis* in 10% LB medium. Thus, we controlled the population ratio of the two strains (*B. subtilis*/*E. faecalis*) between 0.04 and 0.4%. Therefore, we successfully evaluated the effects of non-dye-decolorizing *B. subtilis* with a 1 % population on the enhancement of the activity of dye-decolorizing *E. faecalis*.

## RESULTS AND DISCUSSION

### Effects of *B. subtilis* on the activity of *E. faecalis*

We investigated the activity of *E. faecalis* T6a1 using the dye decolorization assay at a dye concentration of 60 mg/L. **Fig. 1A** shows typical changes in dye concentrations during the co-incubation of *E. faecalis* T6a1 and *B. subtilis* S4ga and the sole incubation of *E. faecalis* T6a1. The *E. faecalis* T6a1 population during the co-incubation was approximately one-fifth that of the sole incubation at 48 h: 1.8 × 10^8^ CFU/mL for the co-incubation, 8.4 × 10^8^ CFU/mL for the sole incubation. However, the dye concentration decreased earlier and faster during the co-incubation, and at 48 h, the amount of dye decolorized during the co-incubation was almost twice that of the sole incubation. In contrast, the sole incubation did not show additional decolorization after 48 h. In the co-incubation, the population of *B. subtilis* S4ga was 1.4 × 10^5^ CFU/mL, and thus, the population ratio of *B. subtilis* S4ga to *E. faecalis* T6a1 was 0.08%; this low ratio resulted in differences in the dye decolorization activity of *E. faecalis* T6a1. Furthermore, the maximum dye decolorization rate of the co-incubation was twice as high as that of the sole incubation (**Fig. 1B**). We also repeatedly observed that the co-incubation doubled the amount of dye decolorized and halved the lag time to start decreasing the dye concentration compared with those of the sole incubation (**Fig. 1C, D**). Additionally, co-incubation increased the duration of dye decolorization by approximately 2.2 times compared with sole incubation of *E. faecalis* T6a1 (**Fig. 1E**). In other words, dye decolorization by the co-incubated strain started earlier and lasted longer compared with that of the sole incubated strain, resulting in decolorization of a large amount of dye. The population ratio of *B. subtilis* S4ga to *E. faecalis* T6a1 was only 0.1%.

**Figure 1.**
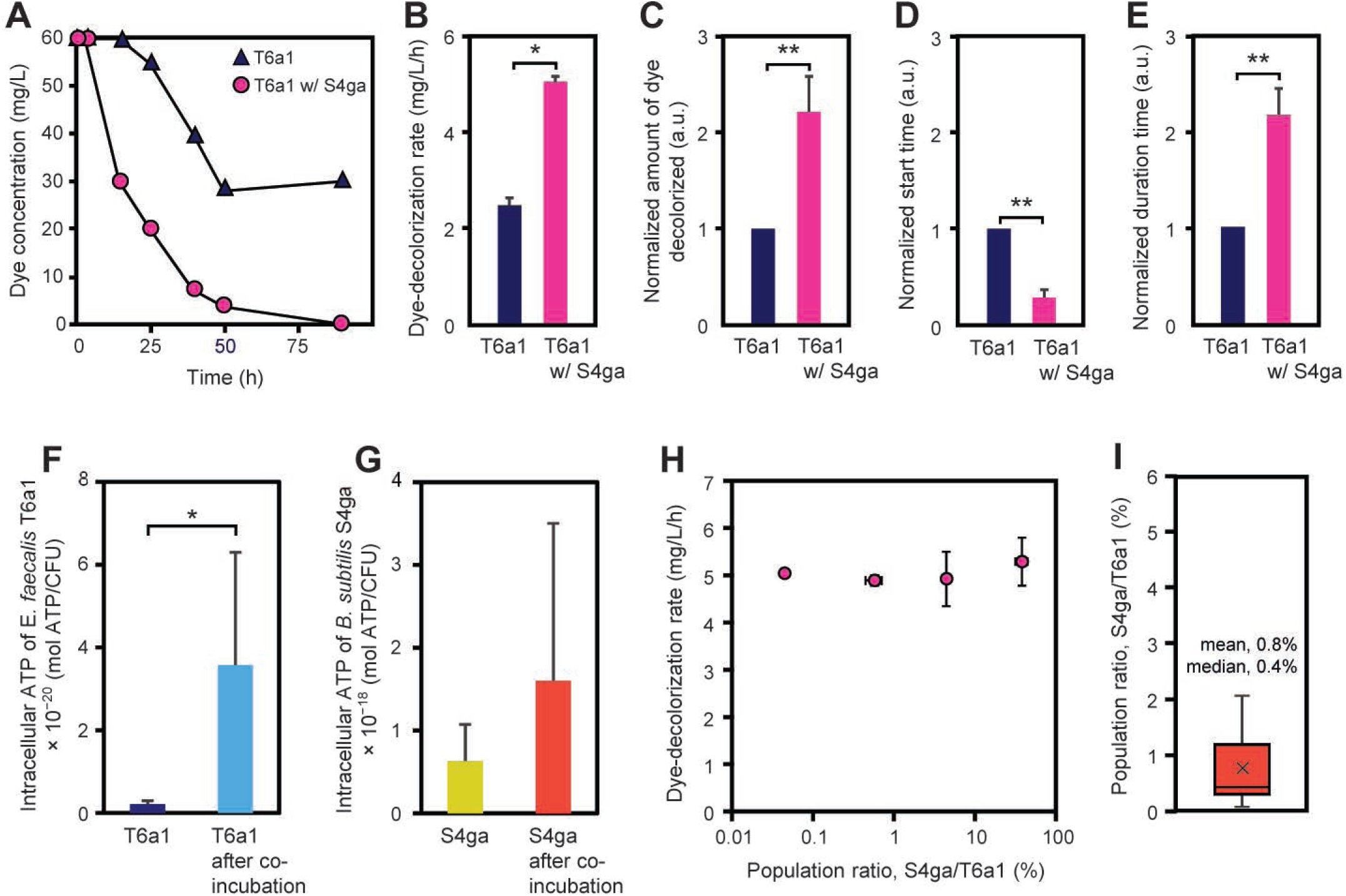
Effects of *B. subtilis* S4ga on dye decolorization of *E. faecalis* T6a1. (A) Dye decolorization over time during co-incubation of *B. subtilis* S4ga and *E. faecalis* T6a1 or sole incubation of *E. faecalis* T6a1. (B) Dye decolorization rates of solely incubated *E. faecalis* T6a1 and co-incubated *E. faecalis* T6a1 with *B. subtilis* S4ga for 11 h. (C) Ratio of dye amount decolorized by the co-incubation to dye amount decolorized by the sole incubation. (D) Ratio of the time when the dye concentration started to decrease during the co-incubation to the time when the dye concentration started decreasing in the sole incubation. (E) Ratio of the time when the dye concentration continued to decrease in the co-incubation to the time when the dye concentration continued to decrease in the sole incubation. (F) Intracellular ATP of solely incubated *E. faecalis* T6a1 or co-incubated *E. faecalis* T6a1 after removal of *B. subtilis* S4ga at 24 h. (G) Intracellular ATP of solely incubated *B. subtilis* S4ga or co-incubated *B. subtilis* S4ga after removal of *E. faecalis* T6a1 at 24 h. (H) Dye decolorization rates of *E. faecalis* T6a1 co-incubated with *B. subtilis* S4ga at the different population ratios ranging from 0.04% to 40%. (I) Distributions of the population ratios of *B. subtilis* S4ga to *E. faecalis* T6a1 that enhanced the dye-decolorization activity of *E. faecalis* T6a1. A box and whisker plot was created with data lower than 4% to focus on the effectiveness of the low ratio range. Error bars represent standard deviations (B– I); \**P* < 0.05; \*\**P* < 0.01; n = 3 (B); n = 9 (C–E); n = 3 (F, G); n = 8 (I). a.u., arbitrary units.

Co-incubation enhanced the decolorization effects of *E. faecalis* T6a1. For instance, shortening the lag time to start decreasing the dye concentration indicated that *B. subtilis* S4ga affected the activity of *E. faecalis* T6a1 before or at the same time that dye degradation proceeded. Therefore, we measured the intracellular ATP levels of co-incubated *E. faecalis* T6a1 and that of the sole incubated *E. faecalis* T6a1. The intracellular ATP level of co-incubated *E. faecalis* T6a1 was 16 times higher than that of the sole-incubated *E. faecalis* T6a1 (**Fig. 1F**). On the other hand, the intracellular ATP of co-incubated *B. subtilis* S4ga was 2.5 times higher than that of the sole-incubated *B. subtilis* S4ga (**Fig. 1G**). These results suggest that both *E. faecalis* T6a1 and *B. subtilis* S4ga were activated by co-incubation. The statistical difference in ATP values in Figure 1G was not sufficiently significant (*P*=0.44), which suggested that the intracellular ATP levels were more variable in the co-incubated *B. subtilis* S4ga, even though the population of *B. subtilis* S4ga during co-incubation was controlled within the same order.

### Population ratio of *B. subtilis* to *E. faecalis*

We evaluated the effect of various population ratios of *B. subtilis* S4ga to *E. faecalis* T6a1 on dye decolorization activity of *E. faecalis* T6a1 by changing the amount of *B. subtilis* S4ga added to the same amount of *E. faecalis* T6a1. The effective population ratios ranged between 0.04% and 40%. There were no significant differences in the decolorization rates of the cocultures within this wide range of *B. subtilis* S4ga population ratios (**Fig. 1H**). However, the decolorization rates of the co-incubations were higher than those of the sole incubation at each time point (**Fig. 1B, H**). We created a box and whisker plot to show the distribution of the effective population ratio with data lower than 4% and demonstrated that 0.4–0.8% *B. subtilis* S4ga (less than 1%) enhanced the dye decolorization activity of *E. faecalis* T6a1 (**Fig. 1I**).

### Effects of *B. subtilis* on the oxidation state

The dye-decolorizing enzyme azoreductase, which is expressed by *E. faecalis*, is activated under reduced environments (**16**). Therefore, we investigated dissolved oxygen concentration (DO) and decolorization rate of cocultures of *E. faecalis* T6a1 and *B. subtilis* S4ga under oxic and anoxic conditions for 23 h (**Fig. 2**). Sole incubation of each strain was performed for comparison. The sole incubation of *E. faecalis* T6a1 showed approximately 75% decolorization under anoxic conditions at 23 h, while the sole incubation of *E. faecalis* T6a1 showed a much lower decolorization rate (20–45%) under oxic conditions at 23 h, mainly between 3.5 and 4.2 mg/L DO (**Fig. 2A**). On the other hand, the co-incubation of *E. faecalis* T6a1 and *B. subtilis* S4ga resulted in higher dye decolorization rates of 80–95% at 8 h under oxic conditions, mainly between 1.5 and 3.2 mg/L DO, and further increased to 90–100% at 23 h. Under anoxic conditions, the co-incubation showed incomplete decolorization of approximately 75–90% at 23 h (**Fig. 2B**). We found that under oxic conditions, co-incubation of *E. faecalis* T6a1 and *B. subtilis* S4ga lowered DO below 3 mg/L, whereas sole incubation of *E. faecalis* T6a1 did not decrease DO as much.

**Figure 2.**
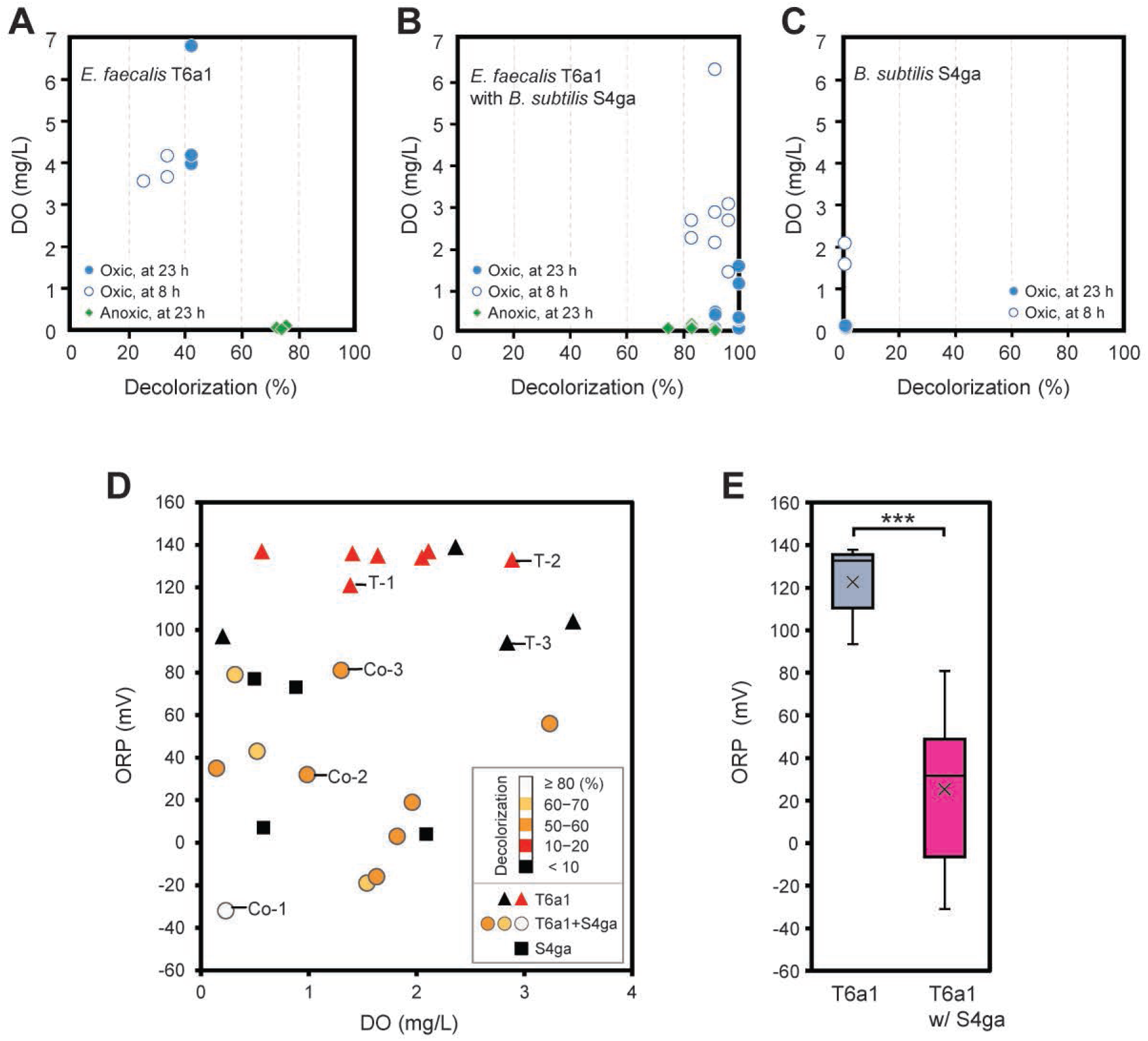
Impact of *B. subtilis* S4ga on dissolved oxygen (DO) concentration and redox potential (ORP) during co-incubation with *E. faecalis* T6a1 and during individual incubation with dye under oxic and anoxic conditions. Correlation between final DO and dye decolorization rates of the incubations of (A) *E. faecalis* T6a1, (B) *E. faecalis* T6a1 with *B. subtilis* S4ga, and (C) *B. subtilis* S4ga, as determined at 8 h and 23 h. (D) Correlation between final ORP and DO values after the incubations of *E. faecalis* T6a, *E. faecalis* T6a1 with *B. subtilis* S4ga, and *B. subtilis* S4ga for 20 h. Populations of *E. faecalis* T6a1 and *B. subtilis* S4ga at 20 h (CFU/mL), Co-1: T6a1, 2 × 10^7^; S4ga, 7 × 10^5^ (S4ga-to-T6a1 population ratio, 3%). Co-2: T6a1, 2 × 10^8^; S4ga, 1 × 10^6^ (S4ga-to-T6a1, 0.6%). Co-3: T6a1, 7 × 10^7^; S4ga, 5 × 10^5^ (S4ga-to-T6a1, 0.7%). T-1: T6a1, 3 × 10^6^T-2: T6a1, 2 × 10^7^. T-3: T6a1, 3 × 10^7^. (E) A box and whisker plot showing ORP values after incubation of *E. faecalis* T6a1 or *E. faecalis* T6a1 with *B. subtilis* S4ga. \*\*\**P* < 0.001; n = 11 (T6a1), n = 11 (T6a1 with S4ga). The initial DO under oxic conditions was between 6 and 7 mg/L (A–E). The initial concentration of Congo red dye was 60 mg/L (A–E).

Next, we examined the effects of *B. subtilis* S4ga on ORP values for dye decolorization of *E. faecalis* T6a1, both sole incubation of *E. faecalis* T6a1 and co-incubation with *B. subtilis* S4ga were carried out under oxic conditions below approximately 3 mg/L DO (**Fig. 2D**). The initial oxidation-reduction potential (ORP) was +163 ± 11 mV in the presence of 1.9 mg/L DO. The ORP values of the sole incubation of *E. faecalis* T6a1 were in the range of +90 mV to +140 mV at 20 h. In contrast, the ORP values of the co-incubation were in the range of −30 mV to +80 mV (**Fig. 2E**).

To confirm the effects of *B. subtilis* S4ga on lowering ORP, we monitored the changes in ORP during the sole incubation of *B. subtilis* S4ga under oxic and microaerophilic conditions (**Fig. 3**). Under oxic condition, *B. subtilis* S4ga lowered ORP to approximately 0 mV after 8-h incubation after decreasing DO below 3 mg/L. Under microaerophilic condition, *B. subtilis* S4ga lowered ORP to approximately + 60 mV after 8-h incubation with decreasing DO below 1 mg/L. *B. subtilis* S4ga created a moderately reduced environment when DO was present and the ORP level under oxic condition was lower than that under microaerophilic condition. These results demonstrate that *B. subtilis* S4ga lowered the ORP to a condition that enhanced dye decolorization of *E. faecalis* T6a1 under oxic conditions and that oxygen was required to enhance dye decolorization of *E. faecalis* T6a1 when co-incubated with *B. subtilis* S4ga. *B. subtilis* has been widely used to produce amino acids (**17**) and vitamins (**18**). Since oxygen is required for the production of these compounds (**19, 20**), *B. subtilis* probably produced such reducing agents under oxic conditions, resulting in the lower ORP values observed in this study.

**Figure 3.**
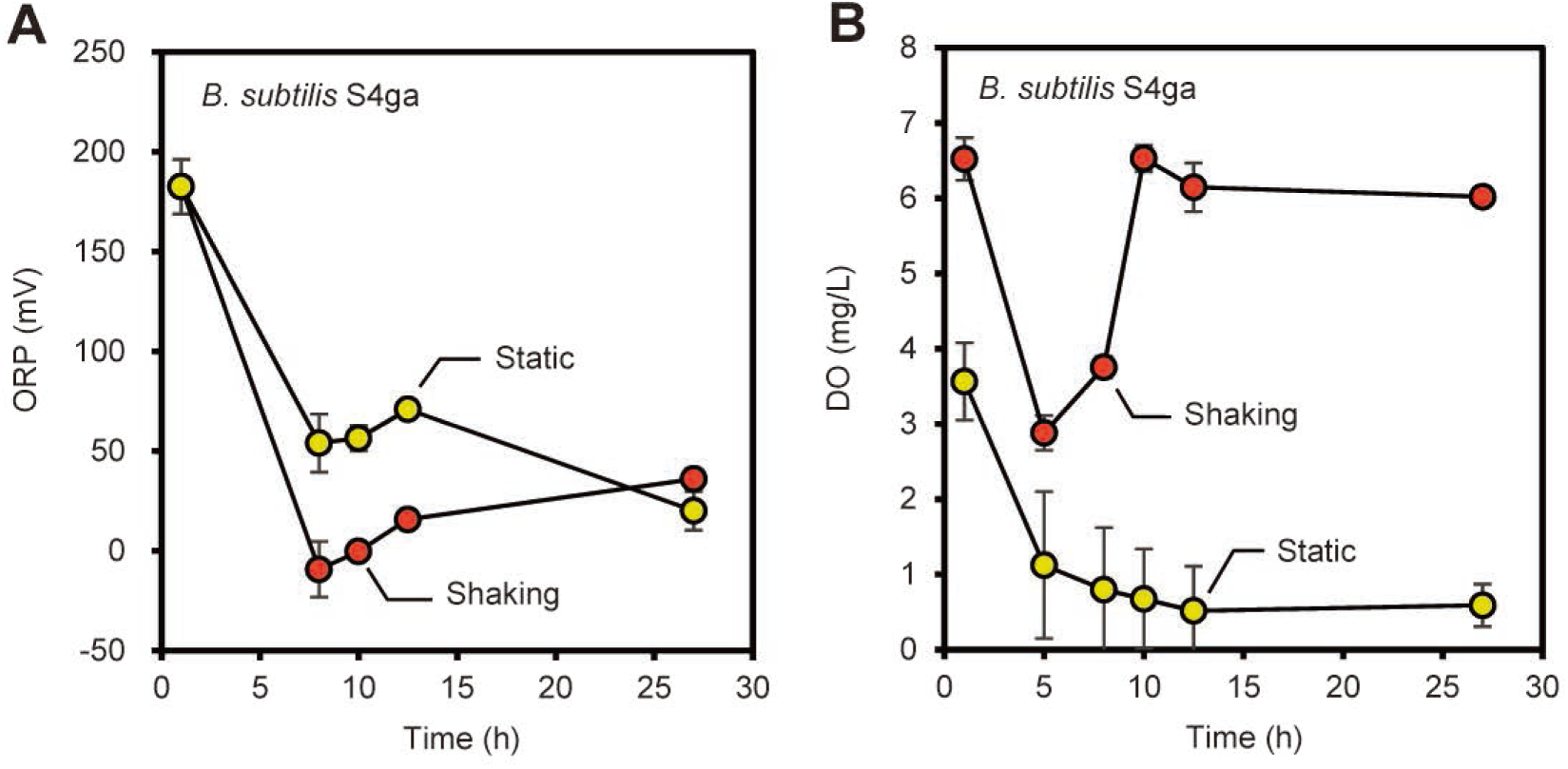
Effects of *B. subtilis* S4ga on redox potential (ORP) under oxic and microaerophilic conditions. ORP (A) and dissolved oxygen (DO) concentration (B) over time during sole incubation of *B. subtilis* S4ga under shaking and static conditions. Rotational shaking at 60 rpm created oxic conditioned. Static incubation after nitrogen purging for nine seconds created microaerophilic condition. Population of *B. subtilis* S4ga at 14 h: oxic, 1 × 10^8^ (CFU/mL); microaerophilic, 0.5 × 10^8^ (CFU/mL). The initial *B. subtilis* S4ga population was 3 × 10^5^ (CFU/mL). Error bars represent standard deviations; n = 3.

### Extracellular metabolites of co-incubation and sole incubations

Extracellular metabolites of individual incubation of *E. faecalis* T6a1 and *B. subtilis* S4ga and co-incubated culture revealed characteristics of amino acid production and consumption from the LB 10% medium (**Fig. 4**). Amino acids produced by *B. subtilis* S4ga were tyrosine, methionine, tryptophan, phenylalanine, valine, and leucine (*P*<0.05). Solely incubated *E. faecalis* T61a utilized all of these amino acids. However, the decreased amounts of methionine, tryptophan, phenylalanine, valine, and leucine in co-culture were not as high as those in the sole incubation of *E. faecalis* T6a1, which is probably because *B. subtilis* S4ga provided the co-culture with these amino acids. Although *B. subtilis* S4ga sole incubation produced tyrosine twice as much as the amount of LB 10% medium, the co-incubation of *E. faecalis* T6a1 with 1% *B. subtilis* S4ga consumed tyrosine and arginine, both of which were possibly rate-limiting substrates. Interestingly, all the amino acids produced by *B. subtilis* S4ga (methionine, tryptophan, phenylalanine, valine, and leucine) were hydrophobic. Since hydrophobic amino acids are more permeable through the cell membrane than hydrophilic amino acids, *E. faecalis* T6a1 can readily acquire these amino acids. In turn, amino acids produced by *E. faecalis* T6a1 were glutamine, histidine, aspartic acid, and proline (*P*<0.05). Solely incubated *B. subtilis* S4ga utilized all of these amino acids. The production and utilization of the amino acids by both strains demonstrated cross-feeding of amino acids between *E. faecalis* T6a1 and *B. subtilis* S4ga, indicating a synergistic effect on the activities of each strain.

**Figure 4.**
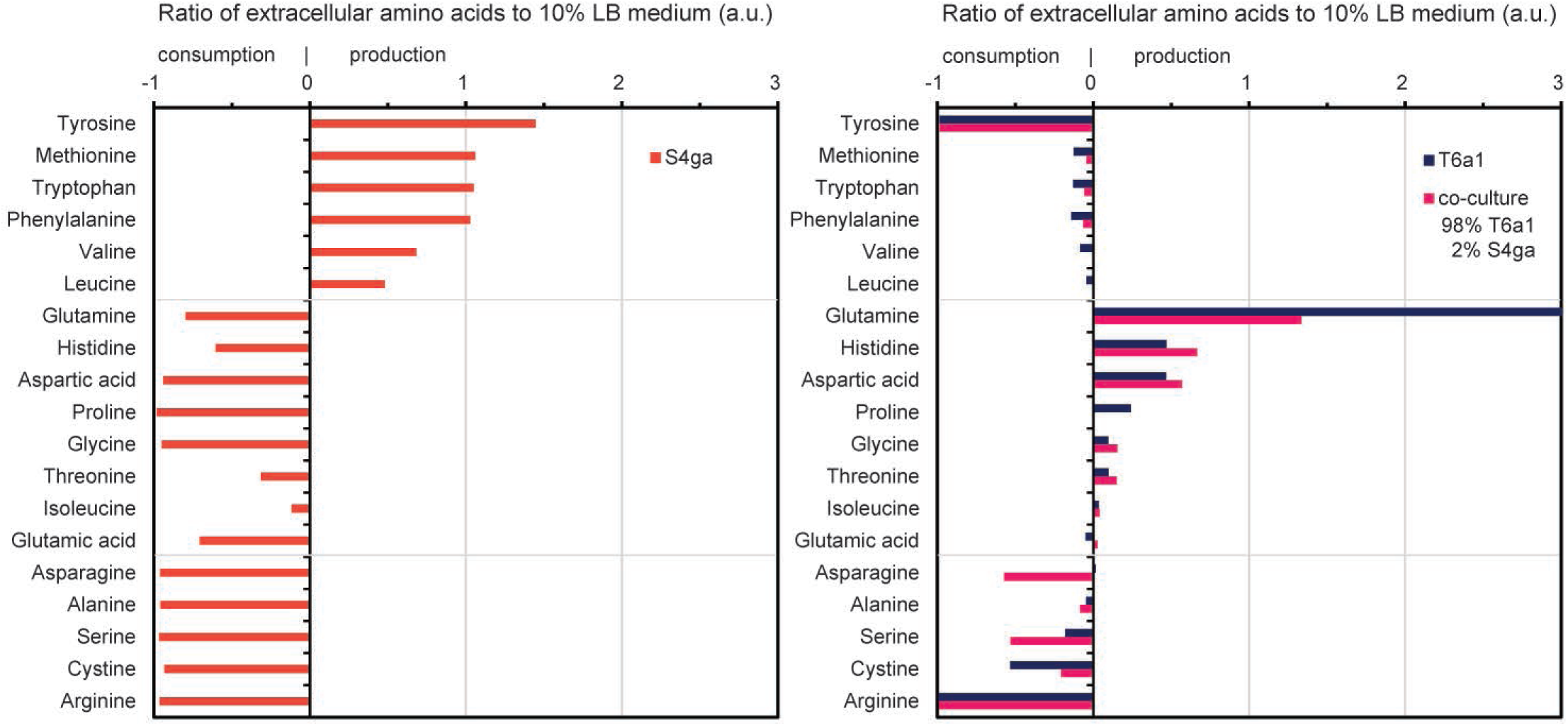
Extracellular amino acids produced or consumed by *B. subtilis* S4ga, *E. faecalis* T6a1 and co-culture of *E. faecalis* T6a1 and *B. subtilis* S4ga. Data represent ratios of amino acids in the supernatants of the incubated cultures at 14 h to amino acids in 10% LB medium at 0 h. Co-culture: *E. faecalis* T6a1, 5 × 10^9^ (CFU/mL); *B. subtilis* S4ga, 1 × 10^8^ (CFU/mL) (S4ga-to-T6a1 population ratio, 2%). n = 3. a.u., arbitrary units.

### Intracellular metabolites of *E. faecalis* incubated with *B. subtilis*

Among the 97 water-soluble primary metabolites, citrulline was the most significantly altered intracellular metabolite, with higher levels in *E. faecalis* T6a1 cells co-incubated with *B. subtilis* S4ga than in solely incubated *E. faecalis* T6a1 cells (**Table S1**). Citrulline is an amino acid produced during arginine degradation by the ADI pathway of *E. faecalis* (**21**). The ADI pathway is an ATP-producing energy-generating process for *E. faecalis* (**22–24**). The ADI pathway catalyzes the conversion of arginine to citrulline and ammonia and further to ornithine, carbon dioxide, and ammonia, with the generation of 2 mol ATP for 1 mol arginine (EC 3.5.3.6, EC 6.3.4.16). In fact, we demonstrated more than 10 times higher amounts of ATP in the co-incubated *E. faecalis* T6a1 cells than in the solely incubated *E. faecalis* T6a1 cells (**Fig. 1F**). Although low oxygen tension is required to induce the ADI pathway in *Pseudomonas aeruginosa* (**25**), in *E. faecalis*, oxygen tension has a weak effect on the ADI pathway enzymes, and arginine is a more important inducer (**23, 24**). Similarly, higher dye-decolorization activities of *E. faecalis* T6a1 were obtained under oxic conditions (**Fig. 2B**), and extracellular arginine was extensively consumed in both solely incubated *E. faecalis* T6a1 and co-incubated *E. faecalis* T6a1 (**Fig. 4**). In addition, among the intracellular metabolites of *E. faecalis* T6a1, the detected amount of arginine was higher in the co-incubation than in the sole incubation of *E. faecalis* T6a1 (**Fig. 5, Table S1**). Together with citrulline and arginine, the increase in ornithine is also explained by the observation that the ADI pathway excretes abundant ornithine (**21**). Accordingly, we concluded that co-incubation with *B. subtilis* S4ga activated the ADI pathway and promoted energy generation in *E. faecalis* T6a1 cells.

**Figure 5.**
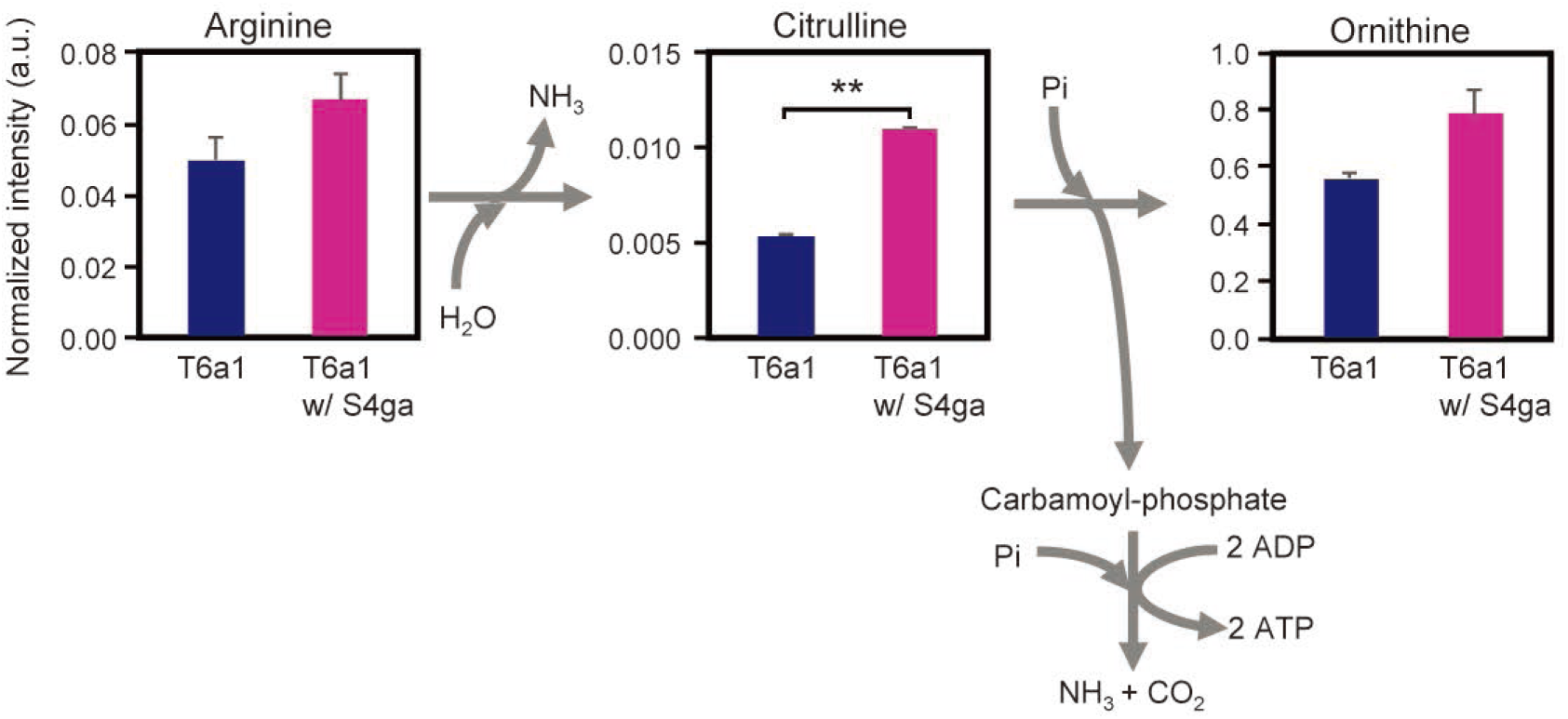
Metabolites of the arginine deaminase (ADI) pathway produced by *E. faecalis* T6a1 during co-incubation with *B. subtilis* S4ga and dye decolorization. Error bars represent standard deviations; \*\**P* < 0.01; n = 2. Data represent normalized intensities of the metabolites. a.u., arbitrary unit.

Proline levels were also significantly higher in cocultured *E. faecalis* T6a1 cells. Proline is an abundant amino acid in *E. faecalis* enterolysin A (**26**), which breaks down bacterial cell walls. Because some strains of *Bacillus*, including *B. subtilis*, show weak sensitivity to enterolysin A (**26**), the presence of *B. subtilis* S4ga may stimulate enterolysin A production by *E. faecalis* T6a1. Moreover, we assumed that the presence of *B. subtilis* S4ga affected the oxidative stress response of *E. faecalis* T6a1 and therefore focused on metabolites related to oxidative stress responses, such as glutathione and oxidized glutathione (**27, 28**). It seems that the amount of glutathione detected in solely incubated *E. faecalis* T6a1 was higher than that in the co-incubation, although the difference was not statistically significant. Further studies are needed to determine whether the oxidative stress level of *E. faecalis* T6a1 was decreased by co-incubation with *B. subtilis* S4ga.

In conclusion, we report here that *B. subtilis* S4ga enhanced the activity of *E. faecalis* T6a1, even when *B. subtilis* S4ga was present at only 0.1–1% of the total population, as demonstrated by the dye decolorization assay using non-dye-decolorizing *B. subtilis* S4ga and dye-decolorizing *E. faecalis* T6a1. The effects of *B. subtilis* S4ga on increasing the activity of *E. faecalis* T6a1 appeared not only as a high maximum dye decolorization rate, but also as a shortened lag time for *E. faecalis* T6a1 to start decreasing the dye concentration, as an increased duration of dye decolorization, and the amount of dye decolorization was doubled. During co-incubation, *B. subtilis* S4ga lowered the ORP to approximately 0 mV by consuming oxygen and/or producing reducing agents such as amino acids. The amino acids produced by *B. subtilis* S4ga were the same as those consumed by *E. faecalis* T6a1, while the amino acids produced by *E. faecalis* T6a1 were the same as those consumed by *B. subtilis* S4ga. In other words, these strains had a mutual supply relationship with the amino acids. This elevated the intracellular ATP content of *E. faecalis* T6a1 and *B. subtilis* S4ga, partly via activation of the ADI pathway of *E. faecalis* T6a1. In the future, we will clarify whether the combination of these species or the presence of *B. subtilis* is important for enhancing the whole-cell activity of coexisting bacteria.

## MATERIALS AND METHODS

### Strains used in this study and identification

One bacterial strain (*E. faecalis* strain T6a1) with dye-decolorizing ability was isolated from the fingertips, as described by Ito et al. (**29, 30**). Seven non-dye-decolorizing *Bacillus*-like strains (S4ga, S4gb, C16caA, C47ea, A14d3B, D3da, and C8ba) were isolated in the same manner (**Table S2**). Full-length 16S rRNA gene sequences of these strains were analyzed by colony polymerase chain reaction, followed by Sanger sequencing, which were carried out at GENEWIZ Japan Corp. (Tokyo, Japan). Identification of the strains was based on an NCBI BLAST search for 16S rRNA gene sequences, colony morphology, and microscopy.

### Screening of non-decolorizing *Bacillus* strains that enhanced dye decolorization of *E. faecalis* T6a1

The isolated strains were pre-incubated under two conditions: sole incubation of each strain and co-incubation of *E. faecalis* T6a1 with one *Bacillus*-like strain. Each strain was inoculated into 5 mL of LB liquid medium in a test tube with a lid. After culturing statically for 17 h at 37°C, 30 μL of 10 g/L Congo red dye (Wako Pure Chemicals, Tokyo, Japan; cat. no. 032-03922) was added to each test tube to achieve an initial dye concentration of 60 mg/L. After the addition of Congo red dye, the samples were incubated statically at 37°C under oxic conditions by leaving the gas phase of the test tubes as air. The color of the liquid in the test tubes was regularly observed. The decreased dye concentration was determined by measuring the absorbance of the liquid in the test tubes at a wavelength of 490 nm. When measuring the absorbance, the liquid containing the cells and dye was centrifuged at 14000 × *g* for 5 min, and the supernatant was analyzed using a spectrophotometer. The dye decolorization rate was calculated from the maximum slope of the line on each dye concentration-time graph. The dye decolorization assay was performed in triplicate.

Among the seven *Bacillus* strains, strains S4ga and S4gb were closely related to *B. subtilis* (100% similarity for the full-length 16S rRNA gene). In addition, strain S4ga showed the highest acceleration of dye decolorization of *E. faecalis* T6a1, resulting in complete decolorization of 60 mg/L Congo red to 0 mg/L (**Table S2**). Sole incubation of *E. faecalis* T6a1 decreased the dye concentration, but at a decolorization rate of two-thirds that of the co-incubation with strain S4ga. Thus, strain S4ga was selected for subsequent analyses.

### Population ratio setting

To test different population ratios of *E. faecalis* T6a1 and *B. subtilis* S4ga, we prepared pre-incubated *E. faecalis* T6a1 cells that were suspended in 10% LB medium with high turbidity and pre-incubated *B. subtilis* S4ga cells as follows. *E. faecalis* T6a1 and *B. subtilis* S4ga were separately pre-incubated in 100% LB medium at 37°C for 14 h. The cells were collected and washed with 10% LB medium, and each strain was then suspended in 10% LB medium. The cell turbidity of *E. faecalis* T6a1 was adjusted to approximately 3 O.D. in 10% LB medium, which suppressed the growth of the strains by 2–8 times compared with their initial cell numbers during subsequent incubation with dye. This enabled the regulation of different population ratios of *B. subtilis* S4ga to *E. faecalis* T6a1 during incubation. The population ratios were set to be approximately 1% and in the range between 0.01% and 50%, as estimated based on the cell turbidity after pre-incubation and confirmed by CFU counting. The cell number of *E. faecalis* T6a1 in sole incubation was adjusted to be similar to that of *E. faecalis* T6a1 in co-incubation with *B. subtilis* S4ga.

### Dye decolorization assay

The dye-decolorizing *E. faecalis* T6a1 was co-incubated with the non-dye-decolorizing *B. subtilis* S4ga and Congo red dye at 37°C. In parallel, *E. faecalis* T6a1 sole incubation was used as a control. Each strain was inoculated into 5 mL of 10% LB liquid medium in a test tube with a lid. To assay dye decolorization under oxic conditions, oxic conditions were assessed by leaving the gas phase as air. The initial DO under oxic conditions was between 6 and 7 mg/L. To assay dye decolorization under anoxic conditions, anoxic conditions were created by nitrogen purging of the gas phase in the tube for 60 s. After nitrogen purging, the DO in the liquid in the test tube was less than 0.5 mg/L. The concentration of Congo red dye was set to 60 mg/L. Immediately after the addition of the dye, the DO was measured routinely using a noncontact-type DO sensor, Fibox3-trace (PreSens Precision Sensing, Regensburg, Germany) until the liquid in the test tube was decolorized or the incubation was terminated. The ORP value of the liquid was measured when the incubation was terminated or when the dye in the liquid was completely decolorized. The initial ORP values were in the range of +155 mV to +175 mV. The pH did not change significantly before and after the incubation, which was between 7.2 and 7.6. The viable cell count as CFU was evaluated after decolorization. The dye concentration during incubation was determined from the RGB of the routinely acquired images (**Fig. S2**). The final dye concentration was determined by measuring the absorbance at 490 nm wavelength.

### Viable cell counts of *E. faecalis* T6a1 and *B. subtilis* S4ga

The amount of dye decolorized in the test tube was almost proportional to the population of dye-decolorizing *E. faecalis* T6a1. Therefore, *E. faecalis* T6a1 cell numbers were enumerated by dilution plating before and after incubation. The incubated solutions collected from sole incubation of *E. faecalis* T6a1 and co-incubation of *E. faecalis* T6a1 and *B. subtilis* S4ga were spread on 10% LB agar medium with serial dilutions and then incubated at 37°C for 12 h, followed by colony counting. Although *B. subtilis* S4ga colonies appeared with *E. faecalis* T6a1 colonies on the same agar plate, they were easily discriminated based on their colony sizes (**Fig. S1**). *E. faecalis* T6a1 lacked motility and was obviously smaller in size (colonies measured approximately 0.5 mm in diameter), whereas *B. subtilis* S4ga was motile and had larger colony sizes (1.5–3 mm in diameter) after incubation for 12 h. *B. subtilis* S4ga colonies were further confirmed by their *Bacillus*-like wrinkled appearance after incubation for 24–48 h.

### Dye concentration estimation by measuring RGB values in photos of incubating tubes

Time-course monitoring of dye concentration was performed by measuring the RGB values in the photographs of the test tubes during incubation. First, standard solutions of Congo red dye were prepared in 10% LB medium at concentrations ranging from 0 to 300 mg/L to obtain a calibration equation. Five milliliters of each standard solution was placed in a flat-bottom test tube for spectrophotometer analysis (Spectroquant Empty cells 16 mm; Merck KGaA, Darmstadt, Germany), which was also used for incubation. All test tubes of the standard solutions were placed in a photography box equipped with an LED backlight, and photographs of the test tubes were obtained using a camera with the following settings: shutter speed, 1/100 s; aperture, f/2.8; focal length, 4.5 mm; ISO speed, 100 (**Fig. S2A**). After loading the images on a PC, the values of R, G, and B at the center of each test tube were measured using the Eyedropper tool in PowerPoint (Microsoft Corporation, Redmond, WA, USA). The total RGB value was calculated for each standard solution to create calibration curves of the total RGB value versus dye concentration, yielding a linear approximation for concentrations below 40 mg/L and an exponential approximation between 40 and 300 mg/L (**Fig. S2B**). When determining the dye concentration in the incubation tubes, photographs of the tubes were taken along with test tubes of the standard solutions with initial dye concentrations of 60 and 300 mg/L as internal standards.

This method is a non-contact type measurement, which does not require collecting the liquid from the test tube and thus does not disturb the incubation solution in the test tube and the gas phase of the tube. Furthermore, the measurement can be performed quickly and repeatedly by taking photographs of the test tubes.

### ORP measurements during *B. subtilis* S4ga incubation

A part of the *B. subtilis* S4ga colony on an agar plate was suspended in 50 mL of LB liquid medium and placed in an incubator at 37°C for 1 h. The cell suspension was centrifuged at 12000 rpm for 5 min and the supernatant was replaced with 10% LB liquid medium. The centrifuged cells were re-suspended in 10% LB and thus a *B. subtilis* S4ga cell suspension (8 log CFU/mL) was prepared in a 50 mL tube. For aerobic incubation, 200 mL of 10% LB liquid medium was prepared in a 500 mL vial, and 240 μL of *B. subtilis* cell suspension was added to the medium. Congo red (10 g/L) was then added at a concentration of 60 mg/L. Three sets of vials were shaken at 60 rpm. For microaerophilic incubation, 5 mL of 10% LB liquid medium was prepared in a 16 mL test tube with a rubber stopper, and 30 μL of *B. subtilis* S4ga cell suspension was added to the medium. Congo red (10 g/L) was added at a final concentration of 60 mg/L. After the tube was capped with a rubber stopper, nitrogen gas was injected into the test tube with a needle for 3 s. DO was 1 mg/L at 5 h and thereafter remained lower than 1 mg/L. During incubation at 37°C, the ORP and DO of the incubation medium of the vials and test tubes were monitored for 27 h. Since it is necessary to open the caps of the vials and tubes for measuring ORP, DO was measured using a non-contact-type DO sensor before opening them. To measure the ORP of the incubation solution under microaerophilic conditions, unopened test tubes were opened each time. Therefore, 30 test tubes were initially prepared, and four to six test tubes were measured at each time point.

## Metabolomic analysis

### (i) Incubation conditions

To investigate whether *B. subtilis* S4ga had some effects on the metabolism of *E. faecalis* T6a1, metabolomic analysis of the intracellular and extracellular water-soluble primary metabolites of strains T6a1 and S4ga was carried out for coculture and sole incubation cultures of each strain. The incubations were conducted as described above (dye decolorization assay). Briefly, after pre-incubation of each strain, co-incubation of *E. faecalis* T6a1 and *B. subtilis* S4ga and sole incubation of each strain were performed at an initial Congo red dye concentration of 60 mg/L in 10% LB medium at 37°C. The incubations were terminated at 10 h during decolorization. The use of 10% LB medium reduced background amino acids and vitamins derived from tryptone and yeast extract of the incubation medium when intracellular and extracellular metabolites were extracted.

### (ii) Extraction and analysis of intracellular water-soluble primary metabolites of *E. faecalis* T6a1

To compare intracellular metabolites of the co-incubated *E. faecalis* T6a1 with those of the sole incubated *E. faecalis* T6a1, it was necessary to remove *B. subtilis* S4ga cells from the co-incubation solution. To do this, we focused on cell sizes; *E. faecalis* T6a1 cells were spherical, approximately 0.8 μm in width and 1.3 μm in length, whereas *B. subtilis* S4ga cells were elongated, approximately 0.8 μm in width and 2.6 μm or more in length (**Fig. S3A, B**). Therefore, we expected that *B. subtilis* S4ga cells would not pass through a 0.8-μm pore size filter compared with *E. faecalis* T6a1 cells. Accordingly, a 0.8-μm filter was used to reduce the number of *B. subtilis* S4ga cells in the co-incubation solution. After 10-h incubation with dye, the co-incubation solution was filtered through a 0.8-μm pore size membrane filter (DISMIC 25CS080AS; Advantec, Tokyo, Japan). In parallel, the sole incubation solution of *E. faecalis* T6a1 was also filtered to ensure that the filtration procedure did not affect the metabolites in *E. faecalis* T6a1 cells. The filtered solutions were further filtered twice more to minimize contamination by *B. subtilis* S4ga cells. Although *B. subtilis* S4ga accounted for 2% of the total population after the 10-h incubation (**Fig. S3C, D**), the filtration treatment did not allow us to find vegetative cells of *B. subtilis* S4ga, despite observing approximately 10,000 cells of *E. faecalis* T6a1 by microscopy. We observed only some endospores or non-vegetative thin cells (**Fig. S3E, F**), and *B. subtilis* S4ga cells accounted for 0.07% of the total population. Such a small population and the remaining non-vegetative cells would not affect the metabolomic analysis of *E. faecalis* T6a1.

*E. faecalis* T6a1 cells prepared by filtration after co-incubation and sole incubation were washed with phosphate-buffered saline once and then suspended in sterilized ultrapure water. The turbidity of the cell suspension of *E. faecalis* T6a1 under co-incubation and sole incubation was adjusted to be the same among all samples by adding ultrapure water. Methanol was then added to the suspensions at a concentration of 80% to extract the intracellular metabolites. Next, 2-morpholinoethanesulfonic acid (2-MES) and methionine sulfone as internal standards were added to the cell suspensions at a final concentration of 10 μM. After thorough mixing and centrifugation at 20000 × *g* for 5 min, the supernatants of the cell suspensions were collected for metabolomic analysis. Two sets of samples were prepared for *E. faecalis* T6a1 co-incubation and sole incubation.

Ninety-seven water-soluble primary metabolites (e.g., amino acids, organic acids, and coenzymes) were evaluated in intracellular extraction solutions of strain T6a1 by liquid chromatography tandem mass spectrometry (Shimadzu LCMS-8050; Kyoto, Japan) equipped with a pentafluorophenylpropyl column. The resulting intensity was normalized as the height ratio (peak height of each metabolite divided by the peak height of 2-MES). In cases in which a small peak was overlaid with the next peak or in which small peaks were considered noise, such peaks were regarded as undetected metabolites. Metabolic pathways of *E. faecalis* were analyzed using the Kyoto Encyclopedia of Genes and Genomes (KEGG) database (**31**).

### (iii) Extraction and analysis of extracellular water-soluble primary metabolites of co-culture and sole incubation cultures of *E. faecalis* T6a1 and *B. subtilis* S4ga

The incubation solutions of the co-culture and sole incubation cultures of *E. faecalis* T6a1 and *B. subtilis* S4ga were centrifuged at 20000 × *g* for 7 min, and the supernatants were filtered using a 0.2-μm pore size sterile membrane filter (ASFIL 033022SO-SFCA; AS ONE, Osaka, Japan). In addition, 10% LB medium was used as a reference to determine the relative concentration of amino acids from the components of tryptone and yeast extract (**32**). Methanol was then added to the filtered supernatants to a concentration of 80% to extract the extracellular metabolites. Next, 2-MES and methionine sulfone as internal standards were added to the supernatants at a final concentration of 10 μM. Three sets of samples were prepared for the co-culture and sole incubation cultures of *E. faecalis* T6a1 and *B. subtilis* S4ga, and 10% LB medium. In the same manner as for the analysis of intracellular metabolites, ninety-seven water-soluble primary metabolites were evaluated in the extracellular extraction solutions.

### Measurement of intracellular ATP of *E. faecalis* T6a1 and *B. subtilis* S4ga

The amount of intracellular ATP in *E. faecalis* T6a1 in sole culture was determined by subtracting the amount of extracellular ATP of the 0.2 μm filtration permeate from the amount of the total ATP of the culture medium containing cells: ATP_E.faecalis_ = ATP_total_ – ATP_0.2 μm-extracellular_. The amount of intracellular ATP of *B. subtilis* S4ga in sole culture was determined in the same manner: ATP_B.subtilis_ = ATP_total_ – ATP_0.2 μm-extracellular_. For co-culture of *E. faecalis* T6a1 and *B. subtilis* S4ga, the amount of intracellular ATP of *E. faecalis* T6a1 in the co-culture was determined by subtracting the amount of extracellular ATP in the 0.2 μm filtration permeate from the amount of ATP in the 0.8 μm filtration permeate containing extracellular ATP and *E. faecalis* T6a1 cells (but not strain S4ga cells): ATP_E.faecalis_= (ATP_0.8 μm-extracellular_+ ATP_0.8 μm-E.faecalis_) – ATP_0.2 μm-extracellular_.

The amount of intracellular ATP in *B. subtilis* S4ga in the co-culture was determined by subtracting the amount of ATP in the 0.8 μm filtration permeate containing extracellular ATP and *E. faecalis* T6a1 cells (but not *B. subtilis* S4ga cells) from the amount of total ATP in the culture medium containing extracellular ATP and intracellular ATP of both strain cells: ATP_B.subtilis_ = ATP_total_– (ATP_0.8 μm-extracellular_+ ATP_0.8 μm-E.faecalis_). ATP measurement was performed with BacTiter Glo ATP measurement reagent (Promega, WI, USA) and a luminometer (Lu-mini, Viti Life Science Solutions, Shanghai, China).

### Statistical analysis

Statistical significance was determined by two-tailed paired *t-*tests to compare the co-incubation with *B. subtilis* S4ga with the sole-incubations of *E. faecalis* T6a1.

### Data availability

The 16S rRNA gene sequences of the isolated strains are available in the DDBJ/EMBL/GenBank databases under the accession numbers LC557810– LC557817.

## ACKNOWLEDGMENTS

This study was supported in part by JSPS KAKENHI (grant number 15K12369 to T.I.) and by a Grant-in-Aid for JSPS Research Fellow (grant number JP17J11210 to Y.Y.). Metabolomic analysis was performed at the Laboratory for Analytical Instruments, Education and Research Support Center, Gunma University.

