## Supplementary material for "*Bacillus subtilis* with a 1% population enhances the activity of *Enterococcus faecalis* with a 99% population": Fig S1

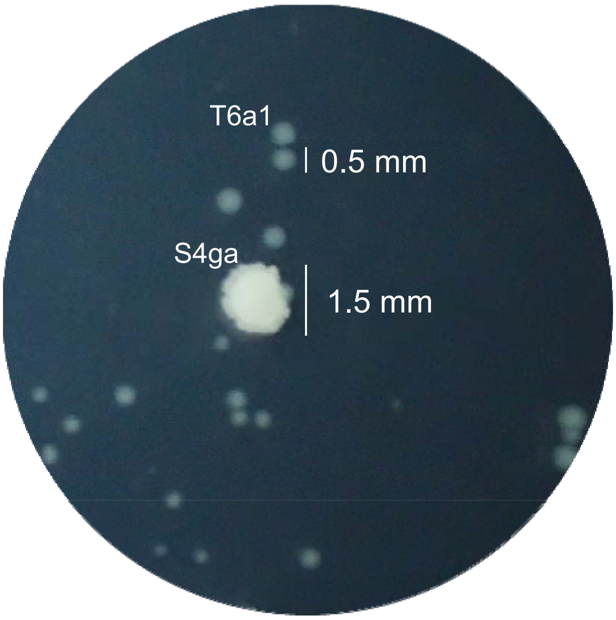


**Figure S1.** Colony images of *E. faecalis* T6a1 (small circular) and *B. subtilis* S4ga (large irregular) on a 10% LB agar plate incubated for 12 h.

**Figure S1.** Ito & Yamanashi
