## Supplementary material for "*Bacillus subtilis* with a 1% population enhances the activity of *Enterococcus faecalis* with a 99% population": Fig S2

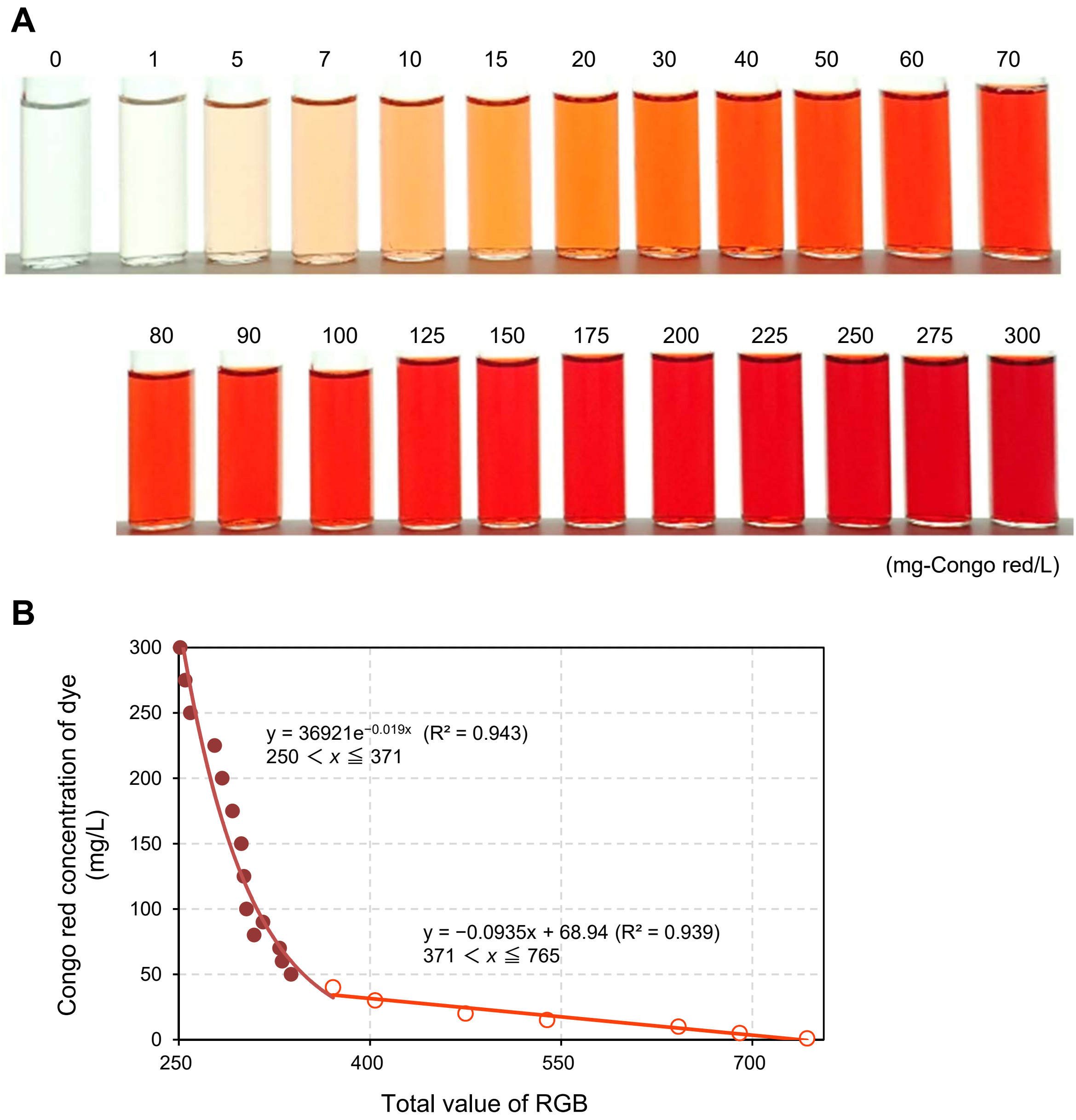


**Figure S2.** Dye concentration estimation by measuring RGB values. (A) Photographs of standard solutions of Congo red dye in 10% LB medium at concentrations ranging from 0 to 300 mg/L. (B) Calibration curve of Congo red concentration created from the total RGB value; linear approximation for concentrations below 40 mg/L and exponential approximation between 40 and 300 mg/L.

**Figure S2.** Ito & Yamanashi
