## Supplementary material for "*Bacillus subtilis* with a 1% population enhances the activity of *Enterococcus faecalis* with a 99% population": Fig S3

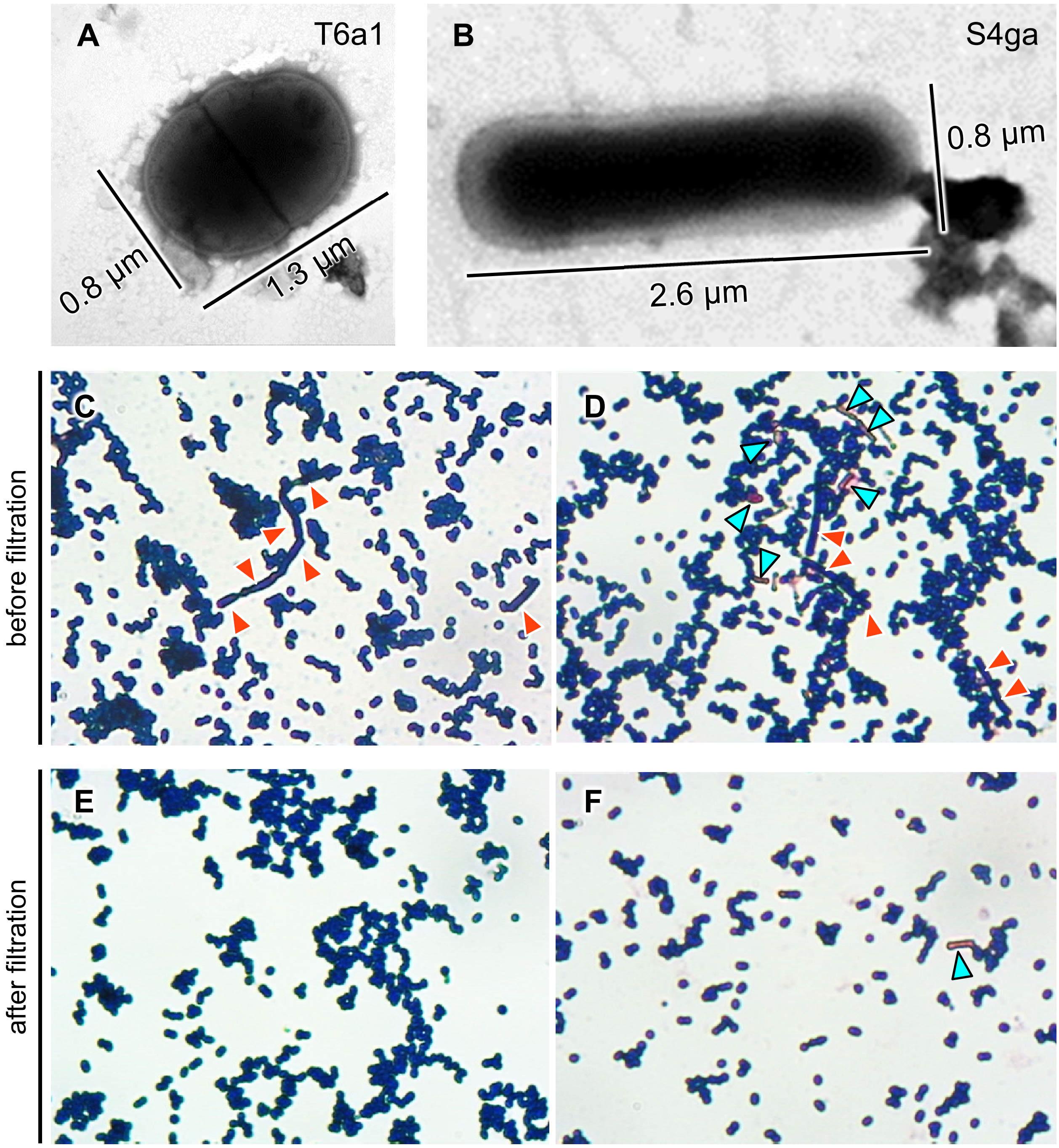


**Figure S3.** Removal of non-dye decolorizing *B. subtilis* S4ga cells from the co-incubation culture with *E. faecalis* T6a1. Electron micrographs of *E. faecalis* T6a1 (A) and *B. subtilis* S4ga (b) when negatively stained with 2% (w/v) phosphotungstic acid and observed under a transmission electron microscope (JEM-1010; JEOL, Tokyo, Japan) at an acceleration voltage of 80 kV. Microscopic images of gram-stained cell suspension of the co-incubation of *E. faecalis* T6a1 and *B. subtilis* S4ga, before (C, D) and after (E, F) 0.8-μm filtration. Both strains were gram-positive (colored blue-violet). Among *B. subtilis* S4ga cells, vegetative cells are indicated with red arrows, whereas endospores and likely nonvegetative thin cells appear pale red (blue arrows).

**Figure S3.** Ito & Yamanashi
