## Supplementary material for "*Bacillus subtilis* with a 1% population enhances the activity of *Enterococcus faecalis* with a 99% population": Table S1

**Table S1.** Normalized intensities of water-soluble primary metabolites extracted from *E. faecalis* T6a1 during dye decolorization when *E. faecalis* T6a1 was co-incubated with *B. subtilis* S4ga and incubated solely.

| Primary metabolite | Normalized intensity | | | | Log2 Fold Change | *p*-value |
|---|---|---|---|---|---|---|
| | T6a1-1 | T6a1-2 | T6a1 w/ S4ga-1 | T6a1 w/ S4ga-2 | | |
| Citrulline | 0.0056 | 0.0052 | 0.0105 | 0.0114 | 1.0168 | 0.007 |
| Adenylsuccinic acid | 0.0023 | 0.0016 | 0.0059 | 0.0044 | 1.4282 | 0.056 |
| Proline | 0.2262 | 0.2584 | 0.4423 | 0.5700 | 1.0628 | 0.057 |
| Histidine | 0.5971 | 0.6319 | 0.9662 | 1.2381 | 0.8428 | 0.071 |
| Ornithine | 0.5348 | 0.5809 | 0.7050 | 0.8674 | 0.4951 | 0.114 |
| S-Adenosylmethionine | 0.0082 | 0.0082 | 0.0109 | 0.0145 | 0.6375 | 0.129 |
| Xanthine | 0.0007 | 0.0004 | 0.0010 | 0.0008 | 0.6977 | 0.139 |
| Alanine | 0.0398 | 0.0405 | 0.0624 | 0.0954 | 0.9734 | 0.143 |
| Adenine | 0.0201 | 0.0173 | 0.0310 | 0.0517 | 1.1466 | 0.162 |
| Uridine | 0.0013 | 0.0009 | 0.0015 | 0.0020 | 0.6938 | 0.167 |
| Aspartic acid | 1.0941 | 1.0808 | 1.3349 | 1.8171 | 0.5353 | 0.180 |
| Hypoxanthine | 0.0019 | 0.0023 | 0.0029 | 0.0048 | 0.8896 | 0.212 |
| Oxidized glutathione | 0.0036 | 0.0062 | 0.0073 | 0.0073 | 0.5721 | 0.215 |
| Arginine | 0.0437 | 0.0560 | 0.0595 | 0.0740 | 0.4207 | 0.218 |
| Threonine | 0.0331 | 0.0391 | 0.0470 | 0.0824 | 0.8422 | 0.253 |
| Methionine | 0.0101 | 0.0087 | 0.0122 | 0.0286 | 1.1196 | 0.311 |
| Guanosine monophosphate | 0.0604 | 0.0592 | 0.0412 | 0.0572 | 0.2808 | 0.320 |
| NAD | 0.0064 | 0.0069 | 0.0069 | 0.0107 | 0.4073 | 0.374 |
| Histamine | 0.0006 | 0.0006 | 0.0009 | 0.0014 | 0.9735 | 0.374 |
| Leucine | 0.0941 | 0.0946 | 0.1040 | 0.3002 | 1.0987 | 0.387 |
| Phenylalanine | 0.0600 | 0.0707 | 0.0679 | 0.2436 | 1.2522 | 0.413 |
| Glutamine | 0.0350 | 0.0647 | 0.0365 | 0.0383 | 0.4149 | 0.490 |
| Isoleucine | 0.0640 | 0.0600 | 0.0784 | 0.1882 | 1.1051 | 0.492 |
| Tyrosine | 0.0134 | 0.0275 | 0.0165 | 0.0454 | 0.5966 | 0.582 |
| Glutathione | 0.0461 | 0.1264 | 0.0472 | 0.0834 | 0.4015 | 0.681 |
| 5-Glutamylcysteine | 0.0751 | 0.1137 | 0.0800 | 0.1372 | 0.2023 | 0.720 |
| Adenosine | 0.2702 | 0.1370 | 0.1535 | 0.2050 | 0.1839 | 0.766 |
| Lactic acid | 0.0746 | 0.1223 | 0.1107 | 0.0996 | 0.0948 | 0.810 |
| Glycine | 0.0058 | 0.0094 | 0.0073 | 0.0088 | 0.0862 | 0.836 |
| 2-Morpholinoethanesulfonic acid[a] | 1.0000 | 1.0000 | 1.0000 | 1.0000 | 0.0000 | - |
| Guanosine | 0.0250 | 0.0179 | 0.0157 | 0.0267 | 0.0163 | - |
| Succinic acid | 0.0211 | 0.0189 | 0.0202 | 0.0208 | 0.0365 | - |
| Niacinamide | 0.0022 | 0.0027 | 0.0023 | 0.0029 | 0.0758 | - |
| Methionine sulfone[a] | 1.8281 | 1.7697 | 1.8318 | 1.5801 | 0.0765 | - |
| Cytidine monophosphate | 0.1923 | 0.1838 | 0.1580 | 0.1961 | 0.0869 | - |
| Serine | 0.0619 | 0.0776 | 0.0732 | 0.0763 | 0.0995 | - |
| Adenosine monophosphate | 0.4267 | 0.4100 | 0.3933 | 0.5678 | 0.1998 | - |
| Acetylcarnitine | 0.0013 | 0.0021 | 0.0017 | 0.0024 | 0.2772 | - |
| Ophthalmic acid | 0.0069 | 0.0088 | 0.0082 | 0.0117 | 0.3454 | - |
| Asparagine | 0.0088 | 0.0151 | 0.0152 | 0.0195 | 0.5437 | - |
| Carnitine | 0.0452 | 0.0383 | 0.0607 | 0.0686 | 0.6321 | - |
| Choline | 0.0131 | 0.0138 | 0.0160 | 0.0279 | 0.7027 | - |
| Cytidine | 0.0265 | 0.0171 | 0.0214 | 0.0511 | 0.7355 | - |
| Methionine sulfoxide | 0.0024 | 0.0021 | 0.0016 | 0.0060 | 0.7488 | - |
| Asymmetric dimethylarginine | 0.0024 | 0.0021 | 0.0030 | 0.0046 | 0.7767 | - |
| Tryptophan | 0.0151 | 0.0158 | 0.0167 | 0.0440 | 0.9742 | - |
| Valine | 0.0757 | 0.0769 | 0.1069 | 0.2375 | 1.1741 | - |
| 4-Aminobutyric acid | 0.0008 | 0.0009 | 0.0014 | 0.0027 | 1.3116 | - |
| Carnosine | 0.0006 | 0.0002 | 0.0007 | 0.0023 | 1.8895 | - |
| Cystathionine | 0.0017 | 0.0069 | 0.0005 | 0.0016 | 2.0563 | - |
| Lysine | 5.4657[b] | 5.4616[b] | 6.9557[b] | 9.7226[b] | 0.6100 | 0.173 |
| Glutamic acid | 2.7319[b] | 2.7431[b] | 3.3127[b] | 4.4373[b] | 0.5013 | 0.180 |

[a] internal standard added.
[b] saturated value.
The metabolites are arranged in order of *p*-value. At the top of the list are significantly different metabolites between the co-incubated and sole-incubated *E. faecalis* T6a1.
Not detected: 2-Aminobutyric acid, Acetylcholine, Aconitic acid, Allantoin, Argininosuccinic acid, cCMP, Cholic acid, Citicoline, Citric acid, Cysteamine, Cystine, Cytosine, Dopa, Epinephrine, Fumaric acid, Homocysteine, Homocystine, Isocitric acid, Kynurenine, Malic acid, Norepinephrine, Orotic acid, S-Adenosylhomocysteine, Serotonin, Taurocholic acid, Thymine, Uric acid, Cysteine.
Weak peak: cAMP, FAD, Guanine, Dopamine, Thymidine monophosphate, Pyruvic acid, Creatinine, cGMP, Nicotinic acid, Inosine, 4-Hydroxyproline, Thymidine, Creatine, Pantothenic acid, Uracil, FMN.
Not determined due to peak overlap: Dimethylglycine, Symmetric dimethylarginine, 2-Ketoglutaric acid.
