## Supplementary material for "*Bacillus subtilis* with a 1% population enhances the activity of *Enterococcus faecalis* with a 99% population": Table S2

**Table S2.** List of non-dye-decolorizing and dye-decolorizing bacterial isolates used in this study and their phylogenetic identification.

| Strain | Dye-decolorization ability | Dye-decolorization rate of co-incubation with strain T6a1 (mg/L/h)[a] | Phylogenetic identification | Sequence similarity (%) | Accession number |
|---|---|---|---|---|---|
| S4ga | − | 5.3 | *Bacillus subtilis* | 100 | LC557810 |
| S4gb | − | 4.9 | *Bacillus subtilis* | 100 | LC557811 |
| C16caA | − | 4.3 | *Bacillus* sp. | 97 | LC557812 |
| C47ea | − | 3.5 | *Bacillus aryabhattai* | 99 | LC557813 |
| A14d3B | − | 3.5 | *Bacillus subtilis* | 99 | LC557814 |
| D3da | − | 3.3 | *Bacillus* sp. | 99 | LC557815 |
| C8ba | − | 3.3 | *Bacillus* sp. | 99 | LC557816 |
| T6a1 | + | 3.5[b] | *Enterococcus faecalis* | 99 | LC557817 |

+; decolorization of dye
−; not decolorized
[a] Incubated in LB medium
[b] Sole incubation of strain T6a1
